# Taking the perspective of an embodied avatar modulates the temporal dynamics of vicarious pain and pleasure: a combined Immersive Virtual Reality and EEG study

**DOI:** 10.1101/2024.06.14.598683

**Authors:** V. Nicolardi, M.P. Lisi, M. Mello, M. Fusaro, G. Tieri, S.M. Aglioti

## Abstract

Observing negative and positive valence virtual stimuli can influence the onlookers’ subjective and brain reactivity. However, the relationship between vicarious pain and pleasure, observer’s perspective taking and cerebral activity remains underexplored. To address this gap, we asked 24 healthy participants to passively observe pleasant, painful, and neutral stimuli delivered to a virtual hand seen from a first-person (1PP) or third-person perspective (3PP) while undergoing time and time-frequency EEG recording. Participants reported a stronger sense of ownership over the virtual hand seen from a 1PP, rated pain and touch valence appropriately, and more intense than the neutral ones. Distinct EEG patterns emerged across early (N2, early posterior negativity, EPN), late (late positive potential, LPP) event-related potentials, and EEG power. The N2 and EPN components showed greater amplitudes for pain and pleasure than neutral stimuli, particularly in 1PP. The LPP component exhibited lower amplitudes for pleasure than pain and neutral stimuli. Further, theta-band power increased, and alpha power decreased for pain and pleasure stimuli viewed from a 1PP versus a 3PP perspective. In the ultra-late time window, we observed decreased theta, alpha, and beta-band power specifically associated with pleasure stimuli. Our study provides novel evidence that perspective-taking modulates the temporal dynamics of vicarious pain and pleasure.

## Introduction

Affective and cognitive variables modulate our first-hand and vicarious sensations (Lisi et al, 2024). While the vicarious experience of others’ feelings and their neural underpinnings has been studied extensively in the domain of pain (Lockwood, 2016; Singer & Lamm, 2009; Betti and Aglioti, 2016) much less attention has been paid to pleasurable touch (Morrison et al., 2011; Lamm et al., 2015; Chiesa et al., 2015, 2017). Interestingly, increasing attention is given to the vicarious experience of pain and pleasure (Pamplona et al., 2022; Mello et al., 2022; Seinfeld et al., 2022) also thanks to the ecological validity of Immersive Virtual Reality (IVR; Tieri et al., 2018). Indeed, IVR typically elicits illusory ownership, i.e. the sensation that the virtual body observed from 1PP is one’s own body (Kilteni et al., 2015). Moreover, seeing virtual pain or pleasure stimuli (e.g. a syringe penetrating or a virtual caress) delivered to an embodied virtual hand elicits in the onlooker unpleasant and pleasurable sensations, respectively (Fusaro et al., 2016, 2019, 2021). However, despite the call for research concerning empathy for positive events (Morelli et al., 2015; Mello et al., 2024) research about the brain correlates of vicarious pleasure mapping is generally limited and almost absent in IVR. In a similar vein, EEG studies about vicarious experience show modulation of components like N1/N2/P3 (Coll, 2018), Early Posterior Negativity (EPN; Fabi and Leuthold, 2017), and Late Positive Potential (LPP; Fan and Han, 2008), as well as modulation of theta and alpha (Mu et al., 2008; Fabi and Leuthold, 2017) and beta responses (Riečanský et al., 2014). However, much less is known about whether vicarious social touch modulates EEG activity. Interestingly, Peled-Avron et al. (2016) demonstrate that behavioural and electrocortical reactivity to observed social touch is modulated by levels of empathy. Schirmer & McGlone (2019) found observing affectionate touch elicited larger LPPs, an electrocortical marker of socio-affective processes but they could not clarify the overlap in brain responses to vicarious pain and pleasure. To fill this gap, we combined an adapted version of our previous IVR paradigm (Fusaro et al., 2016, 2019) with time and time-frequency domain EEG recordings in healthy participants who observed virtual painful, pleasurable, and neutral stimuli, delivered to the hand of an avatar seen from a 1PP (“own” hand) or 3PP (other’s hand). Based on the notion that affective salience, arousal, and attentional engagement are at play in our task, we analyzed EEG in time and time-frequency domain electrocortical components that are known to be involved in sensorimotor and higher-order processes. In particular, we focused on the N2-Visual Evoked Potential, that index selective attention and stimulus intrinsic relevance (Olofsson, 2008; Luck and Kappenman, 2013; Valentini et al., 2017a); the LPP, reflected the arousal and the motivational salience of the stimulus (Cuthbert et al., 2000; Luck and Kappenman, 2011; Hajcak et al., 2009; Valentini et al., 2015), and the EPN that reflects automatic emotion-related influences on information processing (e.g., Olofsson et al., 2008; Schupp et al., 2004). We also focused on: fronto-central theta and centro-lateral alpha power that seem modulated by vicarious experience (Sarlo et al., 2005; González-Franco et al., 2014; Fabi and Leuthold, 2017); fronto-central theta rhythm (Moreau et al., 2020; Valentini et al., 2017a; Cavanagh et al., 2012) that is enhanced by observation of pain and affective touch compared to control conditions (Mu et al., 2008). Finally, we computed central alpha/mu power that appears to decrease when observing painful situations (Mu et al., 2008; Riečanský et al., 2014).

We expected participants to exhibit greater ownership in 1PP vs. 3PP and provide appropriate valence ratings (i.e., pleasant to touch, unpleasant to pain, and neutral to control clip). Furthermore, we expected N2-VEP, LPP and EPN to have greater amplitude associated to vicarious pain and pleasure, compared to neutral, and in 1PP compared to 3PP. We also expected i) stimulus valence and perspective affect fronto-central theta and central alpha power; ii) theta power increases for pain and pleasure compared to neutral, and in 1PP vs 3PP; alpha power decreases for pain and pleasure compared to neutral, and for 1PP vs 3PP.

## Materials and methods

### Participants

Twenty-four healthy volunteers took part in the study (nine males; age mean ± SD, 27.25 ± 3.95). All participants were right-handed with normal visual acuity and were naive as to the purposes of the experiment. The experimental protocol was approved by the ethics committee of the IRCCS Santa Lucia Foundation and was carried out in accordance with the ethical standards of the 2013 Declaration of Helsinki. All participants gave their written informed consent to take part in the study. The number of participants was determined by a priori power analysis performed with MorePower 6.0 (Campbell, 2012) which indicated that 24 participants would be required to detect an effect size of ƞ2p = 0.19 with a power of 0.80 and an α level of 0.05. This estimation was based on previous studies by our group using the same experimental paradigm (Fusaro et al., 2016, 2019).

### Experimental stimuli and setup

The virtual environment was designed using 3DS Max 2017 (Autodesk, Inc.) and implemented in Unity 2017 game software environment (https://unity.com/). It consisted of a real-size living room equipped with some furniture, lamps and a table where two virtual avatars sat facing each other. A virtual gray panel was placed between the avatars to avoid their reciprocal face view and to leave visible only the bodies (see Fig 1. Panel A). Avatars were created by using the Morph Character System (MCS) Male avatar asset and were presented in Unity. Participants observed the virtual scenario and the virtual upper limb from a first-person perspective (1PP) by means of an Oculus Rift head-mounted display (HMD; www.meta.com), having 110° field of view (diagonal FOV) with a resolution of 2.160 × 1.200. (see Fig 1.A Right Panel).

**Figure 1.**
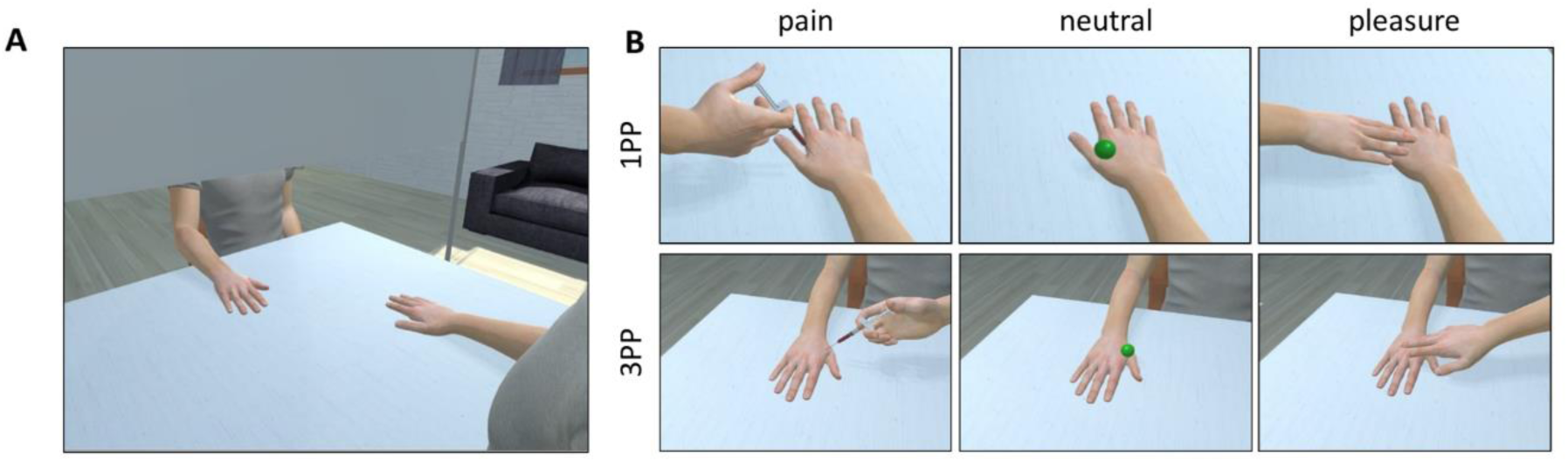
A: representation of virtual environment and the two avatars sat in front of the table facing each other with a grey panel placed between them. B: experimental stimuli observed from 1PP and 3PP (pain, neutral and pleasure).

To reach the best spatial match between real and virtual environment, the virtual table was designed with the same height of the real one and virtual body observed in 1PP was designed and programmed with same posture and position of the participant’s real body. This method, widely used in our previous studies (Tieri et al., 2015a, 2015b, 2017; Fusaro et al., 2016, 2019. 2021; Monti et al., 2020; Fusco et al., 2020), allowed us to implement the basic condition (based on visuo-proprioceptive congruence between the hidden real and observed virtual body) to induce illusory ownership over the virtual body. Experimental stimuli consisted of about three different animations implemented in the virtual environment, with the resulting clips having the same length: i.e. (1) a virtual arm holding a needle on the avatar’s right hand and penetrating it, (2) a virtual hand approaching the avatar’s right hand and caressing it and (3) a virtual green ball approaching and touching the avatar’s right hand (Fig 1, Panel B). The virtual caress was realized in Unity by readapting the real actor’s kinematics used in our previous study (Fusaro et al., 2019). Each stimulus was delivered on the virtual hand observed from 1PP (and thus perceived as ‘own’) or observed from 3PP (and thus perceived as ‘other’). Subjective ratings, concerning the valence of the stimuli (Un/pleasantness ratings), their intensity and the illusory ownership over the observed virtual hand were collected by means of a PC keyboard placed in front of the participant (not displayed in the virtual environment) and interfaced with Unity with a customized C# script (see paragraph *Subjective ratings* for more details). A Neuroscan SynAmps RT amplifiers system (Compumedics) with a 64-channel EEG cap (Electro-Cap International) has been used to record EEG signals (see paragraph *EEG recording* for more details). The EEG system was interfaced with VR by means of a TriggerStation TS832U (Braintrends ltd, www.braintrends.it).

### General procedure

Participants were seated comfortably on a chair in front of a table and wore a 64-channel EEG cap and the HMD (through which they passively observed the virtual body and the scenario from 1PP). Before the experiment, participants underwent the Calibration and Familiarization phases. In the Calibration phase, the point-of-view of each participant were adjusted to pair the 1PP avatar with individual positioning in order to obtain the best spatial-match between the participant’s real and virtual body. In the Familiarization phase, participants (1) were invited to look both at their virtual body and at the environment and to verbally describe what they were seeing (60 sec; Tieri et al., 2015b; Slater et al., 2010) and (2) familiarized with each clip (pain, neutral and pleasure in both 1PP and 3PP) and requested to rate the stimuli-related sensations on the Visual Analogue Scale (VAS). The experiment was composed of two separate blocks (one for each perspective 1PP-3PP) presented in a counterbalanced order across subjects. Each block consisted of 90 trials including each stimulus valence (30 Pain, 30 Pleasure and 30 Neutral presented in pseudo-randomized order). Thus, participants performed a total of 180 trials (90 1PP, 90 3PP). Each trial started with the observation of a right virtual hand (in 1PP or 3PP). After 5500 ± 500 msec, the stimulus was delivered to the virtual right hand on a region overlying the first dorsal interosseous. Each stimulus lasted ∼2.500 msec. Upon its disappearance, participants continued to observe the virtual hand until a sequence of two VAS was presented. Participants were requested to provide ratings about stimulus-related sensations concerning the perceived valence and intensity (see Fig. 2 for the sequence of events per trial). Moreover, three additional VAS were presented to let participants to rate the illusory feeling of ownership over the observed virtual hand (see Table 1, Items 3-5). In particular, in six trials of each block, the Ownership ratings’ VASs appeared in counterbalanced order across trials. Thus, while VAS items 1-2 (Table 1) appeared after each stimulus, VAS items 3-5 were presented in counterbalanced order, 6 times per block, i.e. after 15 trials (two times for pain, pleasure and neutral stimuli in each block).

**Figure 2.**
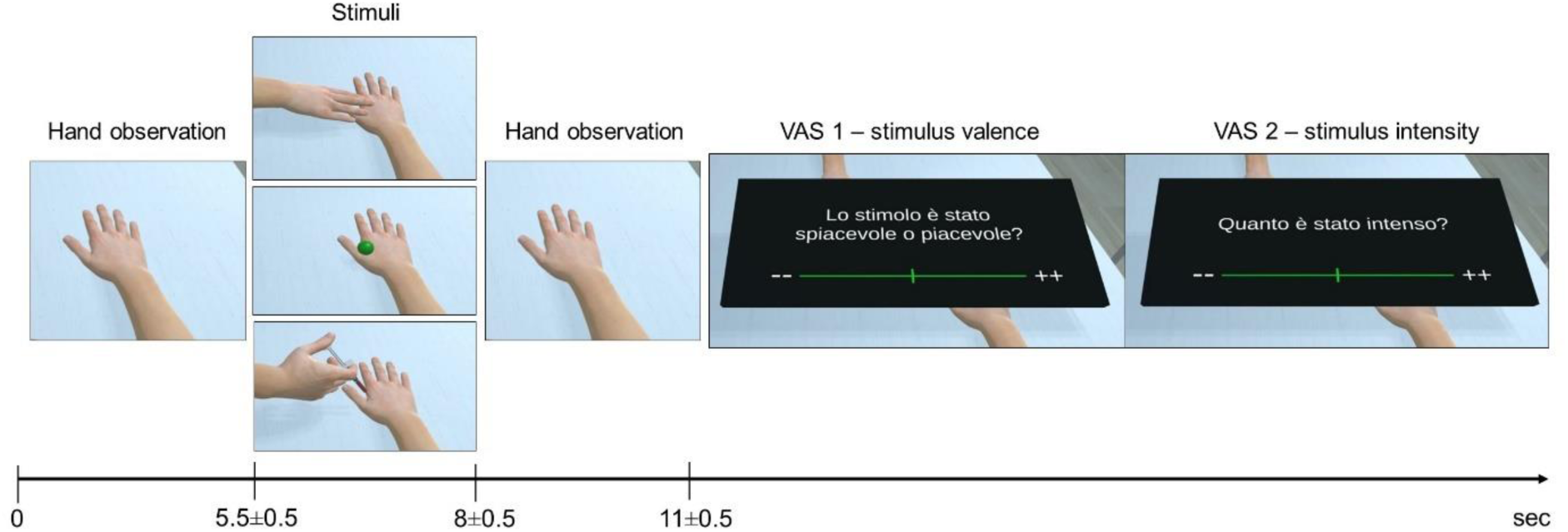
Sequence of events for each trial. VAS 1 and 2, refers to items 1 and 2 of the Table 1.

**Table 1.**
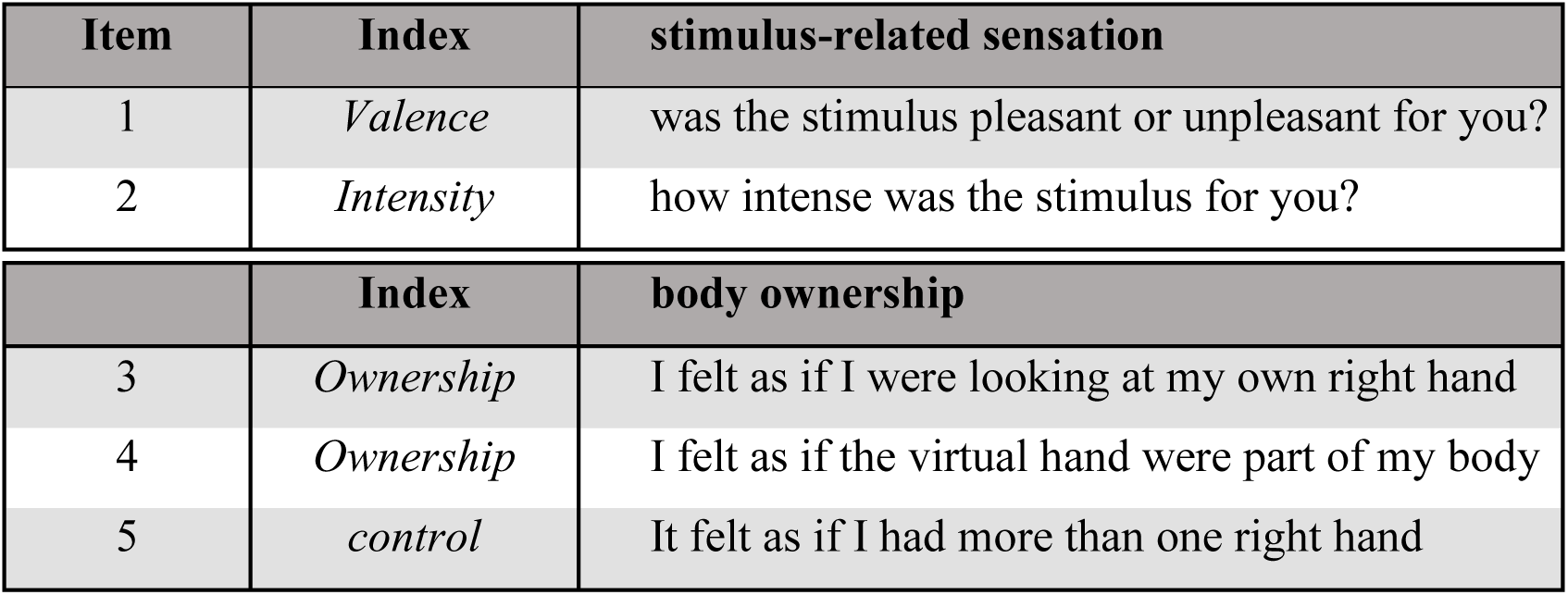
VAS questions concerning the stimulus’s valence (Item 1: “−−” = strongly unpleasant, middle=neutral, “++” strongly pleasant) and intensity (Item 2: “−−” = very weak, “++” = very strong) and VAS statements assessing the illusory feeling of body ownership (Item 3-5: “−−”=no illusion; “++”= maximal illusion).

| Item | Index | stimulus-related sensation |
| --- | --- | --- |
| 1 | *Valence* | was the stimulus pleasant or unpleasant for you? |
| 2 | *Intensity* | how intense was the stimulus for you? |
|  | Index | body ownership |
| 3 | *Ownership* | I felt as if I were looking at my own right hand |
| 4 | *Ownership* | I felt as if the virtual hand were part of my body |
| 5 | *control* | It felt as if I had more than one right hand |

### Subjective Ratings

After each clip, a black panel with a horizontal green line appeared in the virtual space in front of the participants (see Fig. 2). The horizontal green line was 60 cm long with left and right extremities marked as “−−” and “++”, respectively. Participants used their left index and middle finger to control a green cursor along the horizontal line to provide the evaluations listed in Table 1 (items selected and readapted from previous studies; Fusaro et al., 2016, 2019). In particular, item 1 concerned the valence (“−−” = strongly unpleasant, middle=neutral, “++” strongly pleasant) and item 2 the intensity (“−−” = very weak, “++” = very strong;) of the observed clip (Bufalari et al., 2007). Item 3-5 concerned the illusory feeling of body ownership over the virtual hand (“−−”=no illusion; “++”= maximal illusion).

### EEG recording and processing

EEG signals were recorded and amplified using a Neuroscan SynAmps RT amplifiers system (Compumedics Limited, Melbourne, Australia). These signals were acquired from 60 tin scalp electrodes embedded in a fabric cap (Electro-Cap International, Eaton, OH), arranged according to the 10-10 system. The EEG was recorded from the following channels: Fp1, Fpz, Fp2, AF3, AF4, F7, F5, F3, F1, Fz, F2, F4, F6, F8, FC5, FC3, FC1, FCz, FC2, FC4, FC6, T7, C5, C3, C1, Cz, C2, C4, C6, T8, TP7, CP5, CP3, CP1, CPz, CP2, CP4, CP6, TP8, P7, P5, P3, P1, Pz, P2, P4, P6, P8, PO7, PO3, AF7, POz, AF8, PO4, PO8, O1, Oz, O2, FT7, and FT8. All electrodes were physically referenced to an electrode placed on the right earlobe and were re-referenced off-line to the average of all electrodes. Impedance was kept below 5 KΩ for all electrodes for the whole duration of the experiment, amplifier hardware band-pass filter was 0.01 to 200 Hz and sampling rate was 1000 Hz. The continuous EEG data were pre-processed with EEGLAB (Delorme and Makeig, 2004) and Letswave 6 (www.letswave.org). Data were band-pass filtered from 0.1 to 30 Hz (filter order 4) and re-sampled at 250 Hz. Bad channels have been interpolated when needed, based on the signal of the three closest electrodes. Data were segmented into epochs lasting from 1 s before to 1.5 s after the stimulus onset. Eye movements or eye blinks were removed using an Independent Component Analysis (ICA; Jung et al., 2000). The resultant signal was then baseline corrected to a reference interval from −1 to −0.5 s before the stimulus onset and re-referenced to the average reference.

### EEG analyses in the time-domain

In the time domain, middle and late latency visual evoked potentials (VEPs) and Early posterior negativity (EPN) were analyzed separately. Specifically, we focused on: the N2-P2 waveforms, occurring between 200-300 ms which have been proposed to reflect selective attention processes in a complex visual scene (Olofsson et al., 2008); the Late Positive Potential (LPP), a positive deflection occurring after 300 ms (Hajcak et al., 2009), the magnitude of which has been associated with motivational salience, subjective arousal and more in general, processing of emotional contents (Cuthbert et al., 2000; Luck and Kappenman 2012, Valentini et al., 2017a) and the EPN, a negative deflection occurring between 200-300 ms (Farkas et al., 2020), that has been proposed to reflect “natural selective attention”, and to be more sensitive to affective arousing contents (Schupp et al., 2004; Olofsson et al., 2008). VEPs amplitude was measured at occipital ROI (POz-Oz) (Olofsson et al., 2008; Luck and Kappenman, 2012) while EPN amplitude was measured at left parietal ROI contralaterally to stimulus presentation side(P5-P7-PO7) (Frank & Sabatinelli, 2019; Farkas et al., 2020). To investigate any differences between amplitudes across the experimental conditions, single subject average ERPs amplitude for each condition was entered in whole-waveform 2 (Perspective, 1PP vs 3PP) X 3 (stimulus valence, neutral vs pain vs pleasure) ANOVAs. Post-hoc comparisons were performed using the Letswave 6 software, t-tests with correction for multiple comparisons (Maris and Oostenveld, 2007). This method of correction, based on a cluster-level randomisation, allowed us to control the Type-1 error rate involving multiple comparisons. It consists of the following steps: firstly, selecting the points represented by F/t values lower than P < 0.05 to identify the groups (clusters) of contiguous points that show a significant effect; second, estimate the magnitude of each cluster by calculating the sum of the F/t values comprising each cluster. Afterward, random permutation (500 times) of the differences between data-points/pixels at the cluster level within each individual was used to obtain a reference distribution. This distribution was obtained by randomly swapping the conditions within participants and calculating the maximum cluster-level test statistic. For each cluster, a significance threshold was found around the value Z > 2 standard deviations from the mean. Then, for each cluster, a value corresponding to F/t and P (two-tailed) was obtained and the statistical significance ascribed only to differences lower than 0.05 at the end of the permutation process.

### EEG analysis in the time-frequency domain

A short-term Fast-Fourier transform, as implemented in Letswave 6, has been computed to measure responses power in the time-frequency domain. The following parameters were used: frequency between 1-30 Hz, step of 25 for 30 lines, to compute power for each trial, then averaged. The resultant signals, were displayed as an event-related change of average power (ER%), relative to a reference interval before to the onset of the stimulus (−1 to −0.5 s). We focused on the theta and mu power measured at a central ROI (FCz, Cz, C3) that are consided possible indices in perspective taking, conflict monitoring and action mirroring (Pavone et al, 2016; Petereit et al, 2023; Mu et al., 2008; Riečanský et al., 2014).

To investigate any differences in time-frequency domain between experimental conditions (Neutral-1PP, Neutral-3PP; Pleasure-1PP, Pleasure 3PP; Pain-1PP, Pain-3PP), the single subject average power of each experimental condition underwent ANOVA and t-test in Letswave 6, with correction for multiple comparisons (Maris and Oostenveld, 2007, see previous paragraph for details).

## Results

### Subjective Ratings

A graphical representation of subjective reports analysis is provided in Figure 3. *Valence* ratings (Table 1, Item 1) were not normally distributed (Shapiro–Wilk test for one out of six conditions p< 0.05). Thus, nonparametric analysis including Friedman ANOVA and Wilcoxon test for the within-group effects of valence (pain, neutral, pleasure) and perspective (1PP, 3PP), was used. Since Friedman’s ANOVA only allows a One-Way within-subject analysis, we collapsed perspective and valence into a unique 6-level Condition within-subject factor, to test if there is a general effect of Condition. The Friedman ANOVA revealed a significant effect of Condition (χ2(5) = 89.31, p < 0.0001; Kendall’s W= 0.74; Fig. 3A). Bonferroni-corrected Wilcoxon tests for nine planned comparisons (α level = 0.0055) showed that pain (mean ± SD: 27.63 ± 12.87 for 1PP; 26.20 ± 13.28 for 3PP) elicited stronger unpleasantness than neutral (50.75 ± 2.52 for 1PP; 50.75 ± 2.95 for 3PP) and pleasure (66.96 ± 12.56 for 1PP; 64.21 ± 14.46 for 3PP; note that lower and higher ratings represent unpleasant and pleasant sensations, respectively); moreover, pleasure elicited higher pleasantness than neutral and pain (see Table 2-3 for all comparisons). No effect of perspective was found (all ps > 0.05). These results suggest that pain and pleasure stimuli elicited unpleasant and pleasant vicarious sensations, regardless of the perspective from which the stimuli were observed.

**Figure 3.**
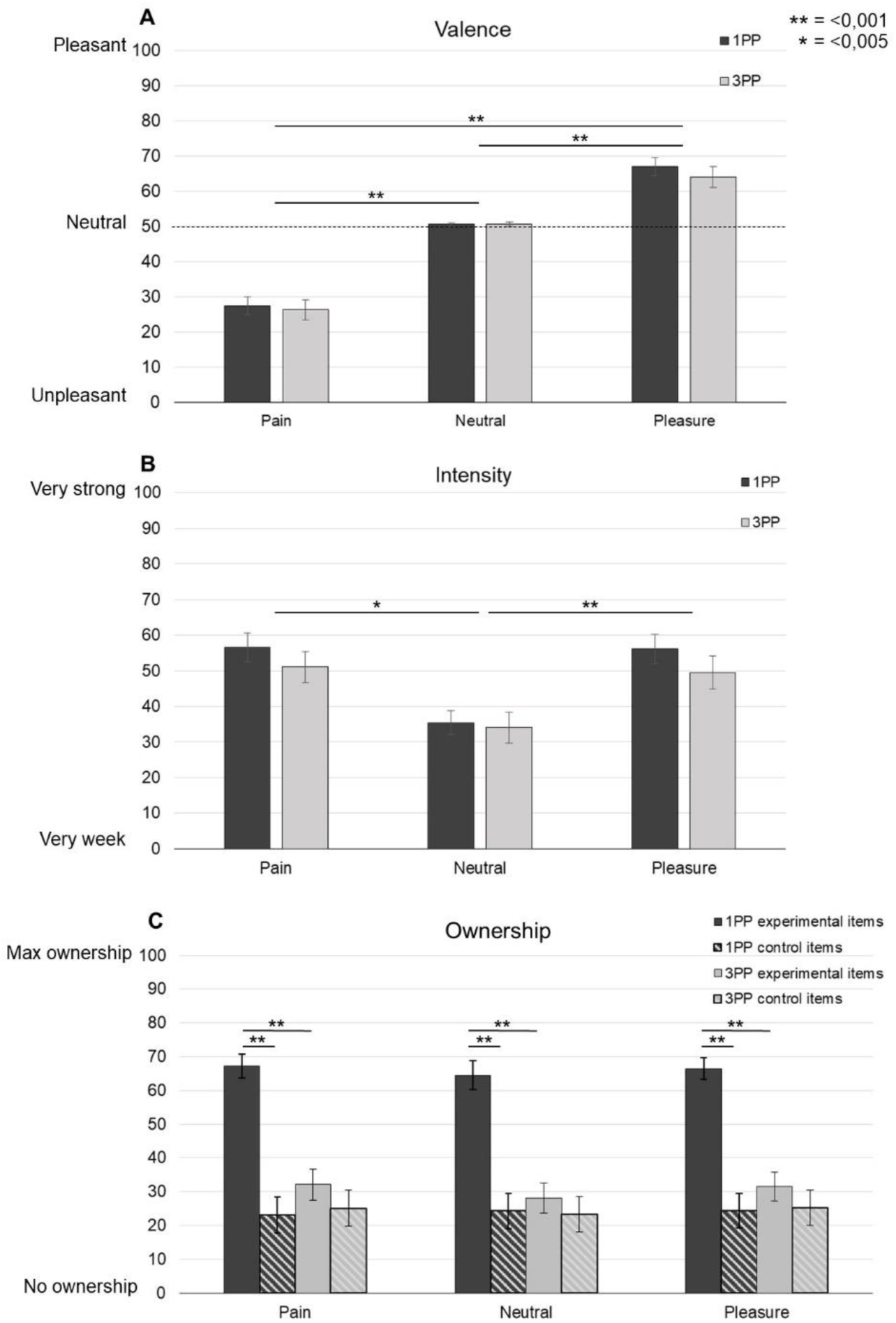
Visual Analog Scale (VAS) ratings (means and SE) of valence (**A**), intensity (**B**), and ownership (**C**) elicited by virtual pain, pleasure and neutral stimuli in 1PP and 3PP.

**Table 2.** Means and SDs of each VAS rating (Valence, Intensity and Ownership) of the type of observed stimulus (pain-pleasure-neutral) in the different Perspectives (1PP and 3PP). High and low UnPleasantness values indicate pleasant and unpleasant sensations, respectively.

|  | First-person perspective (1PP) |  |  | Third-person perspective (3PP) |  |  |
| --- | --- | --- | --- | --- | --- | --- |
| | Pleasure<br>(mean ±<br>SD) | Pain<br>(mean ±<br>SD) | Neutral<br>(mean ±<br>SD) | Pleasure<br>(mean ±<br>SD) | Pain<br>(mean ±<br>SD) | Neutral<br>(mean ±<br>SD) |
| *Valence* | 66.96 ±<br>12.56 | 27.63 ±<br>12.87 | 50.75 ±<br>2.52 | 64.21 ±<br>14.46 | 26.20 ±<br>13.28 | 50.75 ±<br>2.95 |
| *Intensity* | 55.98 ±<br>20.20 | 52.98 ±<br>20.83 | 35.26 ±<br>16.38 | 49.48 ±<br>22.53 | 47.97 ±<br>22.20 | 33.65 ±<br>20.57 |
| *Ownership*<br>*(experimental*<br>*items)* | 67.02 ±<br>15.84 | 68.22 ±<br>17.28 | 64.07 ±<br>21.67 | 32.30 ±<br>22.17 | 33.26 ±<br>24.52 | 28.87 ±<br>23.77 |
| *Ownership*<br>*(control item)* | 23.61 ±<br>22.71 | 23.47 ±<br>23.65 | 25.58 ±<br>24.75 | 2.05 ±<br>24.59 | 23.60 ±<br>24.51 | 22.35 ±<br>25.77 |

**Table 3.** Summary of the main results regarding the subjective Valence, Intensity and Ownership.

| Rating | Post-hoc comparisons (mean ± SD) (* = $p < 0.05$; ** = $p < 0.001$) |
| --- | --- |
| *Valence* | Pain 1PP (27.63 ± 12.87) < Pleasure 1PP (66.96 ± 12.56); n=24; W= 1; r=0.87**<br>Neutral 1PP (50.75 ± 2.52) > Pain 1PP (27.63 ± 12.87) < ; n=24; W= 1; r=0.87 **<br>Neutral 1PP (50.75 ± 2.52) < Pleasure 1PP (66.96 ± 12.56); n=24; W=2; r=0.86 **<br>Pain 3PP (26.20 ± 13.28) < Pleasure 3PP (64.21 ± 14.46); n=24; W=4; r=0.85**<br>Neutral 3PP (50.75 ± 2.95) > Pain 3PP (26.20 ± 13.28); n=24; W=300; r=0.87 **<br>Neutral 3PP (50.75 ± 2.95) > Pleasure 3PP (64.21 ± 14.46); n=24; W=24; r=0.73 ** |
| *Intensity* | Neutral 1PP (35.26 ± 16.38) < Pain 1PP (52.98 ± 20.83); n=24; W= 32; r=0.69 *<br>Neutral 1PP (35.26 ± 16.38) < Pleasure 1PP (55.98 ± 20.20); n=24; W=5; r=0.85 **<br>Neutral 3PP (33.65 ± 20.57) < Pleasure 3PP (49.48 ± 22.53); n=24; W=2; r=0.86** |
| *Ownership* | Control, Pain 1PP (23.47 ± 23.65) < Experimental, Pain 1PP (68.22 ± 17.28); n=24; W= 16; r=0.78**<br>Control, Pleasure 1PP (23.61 ± 22.71) < Experimental, Pleasure 1PP (67.02 ± 15.84); n=24; W= 11; r=0.81**<br>Control, Pleasure 1PP (23.61 ± 22.71) < Experimental, Pleasure 1PP (67.02 ± 15.84); n=24; W= 11; r=0.81**<br>Experimental, Pain 1PP (68.22 ± 17.28) > Experimental, Pain 3PP (33.26 ± 24.52); n=24; W= 290; r=0.82**<br>Experimental, Pleasure 1PP (67.02 ± 15.84) > Experimental, Pleasure 3PP (32.30 ± 22.17); n=24; W= 290; r=0.85**<br>Experimental, Neutral 1PP (64.07 ± 21.67) > Experimental, Neutral 3PP (28.87 ± 23.77); n=24; W= 288; r=0.80** |

*Intensity* ratings (Table 1, Item 2) were not normally distributed (Shapiro–Wilk test for two out of six conditions p < 0.05). We conducted a Friedman ANOVA with the perspective and valence factors collapsed into a unique 6-level Condition within-subject factor. The Friedman ANOVA revealed a significant effect of Condition (χ2(5) = 45.27, p < 0.0001; Kendall’s W= 0.38; Fig. 3B). Bonferroni-corrected Wilcoxon tests for nine planned comparisons (α level = 0.0055) showed significant differences between pain (mean ± SD: 52.98 ± 20.83 for 1PP; 47.97 ± 22.20 for 3PP) vs neutral in 1PP (35.26 ± 16.38 for 1PP; 33.65 ± 20.57 for 3PP) and pleasure (55.98 ± 20.20 for 1PP; 49.48 ± 22.53 for 3PP) vs neutral in both perspectives (see Table 2-3), but not pain vs pleasure (ps > 0.05). No effect of perspective was found (all ps > 0.05). These results suggest that participants perceived as more intense pain than neutral stimuli in 1PP, and pleasure as more intense than the neutral stimuli in both perspectives.

*Ownership* mean scores for experimental (Table 1, Items 3-4) and the control items (Table 1, Item 5) were not normally distributed (Shapiro–Wilk test for six out of twelve conditions p < 0.05). Thus, we conducted a Friedman ANOVA with the item (experimental, control), perspective and valence factors collapsed into a unique 12-level Condition within-subject factor. The Friedman ANOVA revealed a significant effect of Condition (χ2(11) = 100.54, p < 0.0001; Kendall’s W= 0.38; Fig. 3C). Firstly, the mean scores of experimental items and the control one were compared for each condition (1PP-Pain, 1PP-Pleasure, 1PP-Neutral, 3PP-Pain, 3PP-Pleasure, 3PP-Neutral) using Bonferroni correction for six planned comparisons (α level = 0.0083). Wilcoxon tests showed that the scores on the experimental items were higher than the control in 1PP (ps < 0.001), but not in 3PP (ps > 0.05; see

Table 2-3 for descriptive statistics and planned comparisons). Secondly, comparisons showed that this effect was not modulated by valence (α level= 0.0055), as no significant differences were found on the experimental items between 1PP pain (68.22 ± 17.28), 1PP neutral (64.07 ± 21.67) and 1PP pleasure (67.02 ± 15.84; all ps > 0.05, see Table 2-3). These results suggest that illusory ownership was experienced only when the virtual hand was observed from 1PP and regardless of the stimulus valence.

### EEG in the time domain: ERPs results

The ANOVA on VEPs revealed a main effect of valence (pain, neutral, pleasure; F(2,46)=32.83, p<0.001, η^2^p= 0.59) between 160-400 ms, and perspective (1PP, 3PP; F(1,46)=22.17, p<0.001, η^2^p=0.32) between 190-310 ms, on the N2-P2 complex (see Fig. 4). In line with our hypothesis, post-hoc comparisons showed greater amplitude for pain and pleasure as compared to neutral (pain vs neutral: t=−6.53, p<0.001, d=−1.33; pleasure vs neutral: t=−6.55, p<0.001, d=−1.34; Fig.4 panel A) as well as for 1PP compared to 3PP (t=−6.37, p<0.001, d=−1.30; Fig. 4 panel B), for the N2-P2 waveforms. Analysis of LPP showed a main effect of valence (F(2,46)=13.59, p<0.001, η^2^p= 0.23) between 400-600 ms. Surprisingly, this effect was accounted for by greater amplitude for neutral and pain compared to pleasure (pain vs pleasure: t=−4.66, p<0.001, d=−0.95; pleasure vs neutral: t=−4.40, p<0.001, d=0.90; Fig.4 panel A). Figure 4 shows the effect of valence (panel A), and of perspective (panel B); significant post-hoc comparisons are highlighted in the yellow boxes of each panel.

**Figure 4.**
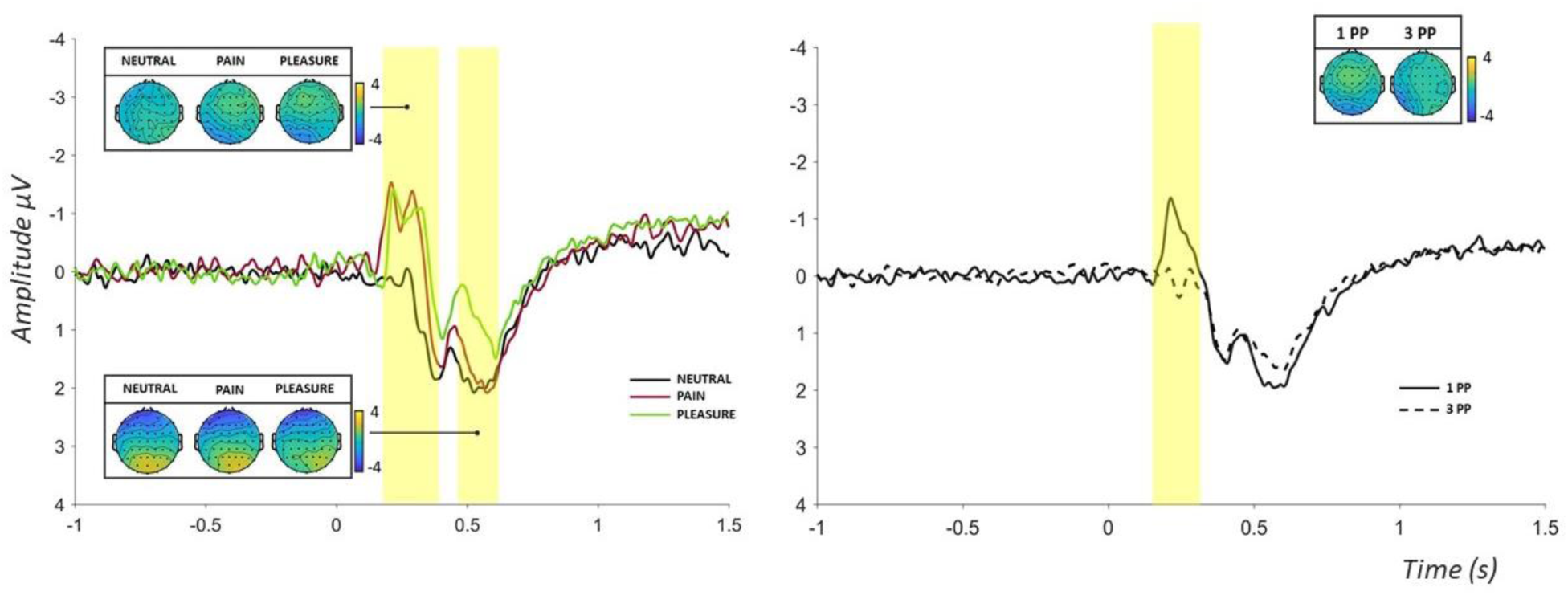
VEPs results. Grand average waveform and scalp topographies of VEPs observed at POz-Oz electrodes. The ANOVA with correction for multiple comparison revealed a main effect of both, valence between 160-400 ms (left, indicated by the first yellow rectangle), and perspective between 190-310 ms (right, in the yellow rectangle), on the N2-P2 complex while a main effect of valence only, on the LPP between 400-600 ms (left, second yellow rectangle), with no significant interaction. Post-hoc comparisons showed Greater amplitude for pain and pleasure as compared to neutral, as well as for 1PP compared to 3PP, was found for the N2-P2 waveforms. The main effect of valence on the LPP was accounted by greater amplitude for neutral and pain compared to pleasure.

The ANOVA on EPN revealed a main effect of valence (F=32.83, p<0.001, η^2^p= 0.59) between 200-535 ms. Post-hoc comparisons indicate greater amplitude for pain and pleasure compared to neutral (pain vs neutral: t=−4.98, p<0.001, d=−1.02; pleasure vs neutral: t=−6.81, p=<0.001, d=−1.39; fig.5, panel A). The main effect of perspective was also significant (F=32.83, p<0.001, η^2^p= 0.42; fig. 5, panel B) between 260-485 ms, an effect that was accounted for by greater amplitude for 1PP compared to 3PP (t=−6.41, p<0.001, d=−1.31). No significant interaction was found.

**Figure 5.**
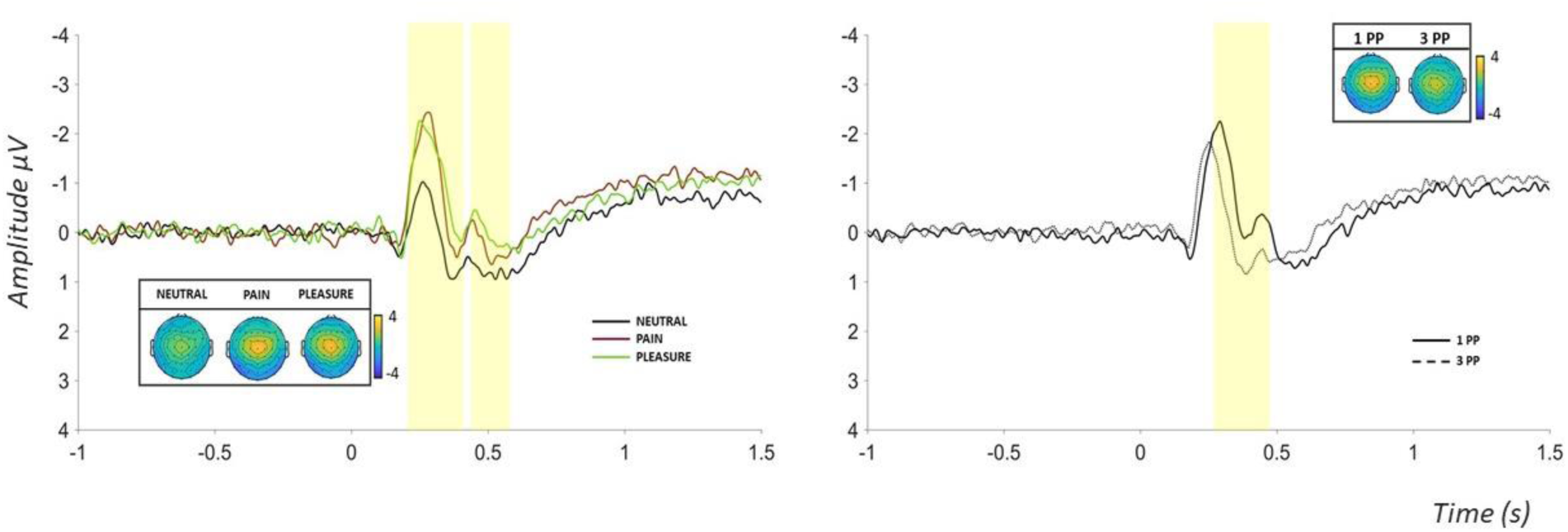
EPN results. Grand average waveforms and topography for EPN at the electrodes P5-P7-PO7. The ANOVA with correction for multiple comparison on EPN revealed a main effect of valence between 200-500ms (left, yellow rectangles) and perspective between 260-500ms (right, yellow rectangle), with no significant interaction. The main effect of perspective was accounted by greater amplitude for 1PP compared to 3PP. Post-hoc comparisons between the levels of valence indicate greater amplitude for pain and pleasure compared to neutral

### EEG in the Time-frequency domain: power changes in different frequency bands

ANOVA on EEG power revealed a main effect of valence (F(2,46)=2.9, p<0.001, η^2^p= 0.11) and perspective (F(1,46)=3.86, p<0.001, η^2^p= 0.08) (see Fig. 6), and a significant interaction between the two (see Fig. 7) (F(2,46)=4.83, p=0.008, η^2^p= 0.17 for the theta range; F(2,46)=10.63, p<0.001, η^2^p=0.32 for the alpha range). Concerning the main effect of valence (Fig. 6, upper row), specific comparisons between the levels of valence (neutral, pain and pleasure), did not show significant differences after correcting p-value for multiple comparisons (500 permutations, see methods). The main effect of perspective (Fig. 6, bottom row), was accounted for by increased theta power in 1PP compared to 3PP in the 300-500ms window (t=3.27, p<0.005, d=0.67), and decreased alpha power in 1PP compared to 3PP in the 500-1000 ms window (t=−2.92, p<0.005, d=−0.59). The significant interaction showed increased theta power in the 200-350ms window, for pain (t=6.08, p<0.001, d=1.24) and pleasure (t=5.29, p<0.001, d=1.08) compared to neutral, in 1PP (Fig. 7, panel A) but not in 3PP (Fig. 7, panel B). Interestingly, in the ultra late time-window (after 800ms), a specific effect of pleasure was found, indicating a power decrease for pleasure compared to neutral. In 1PP, a beta power decrease was found for pleasure compared to neutral (t=−3.01, p=0.006; d=−0.61; Fig. 7, panel A). Moreover, in 3PP, a power decrease for pleasure compared to neutral was found in the theta (t=−3.83, p<0.001, d=−0.78), alpha (t=−3.56, p=0.003, d=−0.73) and beta (t=−3.72, p=0.001, d=−0.76) band (Fig. 7, panel B, second row) in the ultra late time-window.

**Figure 6.**
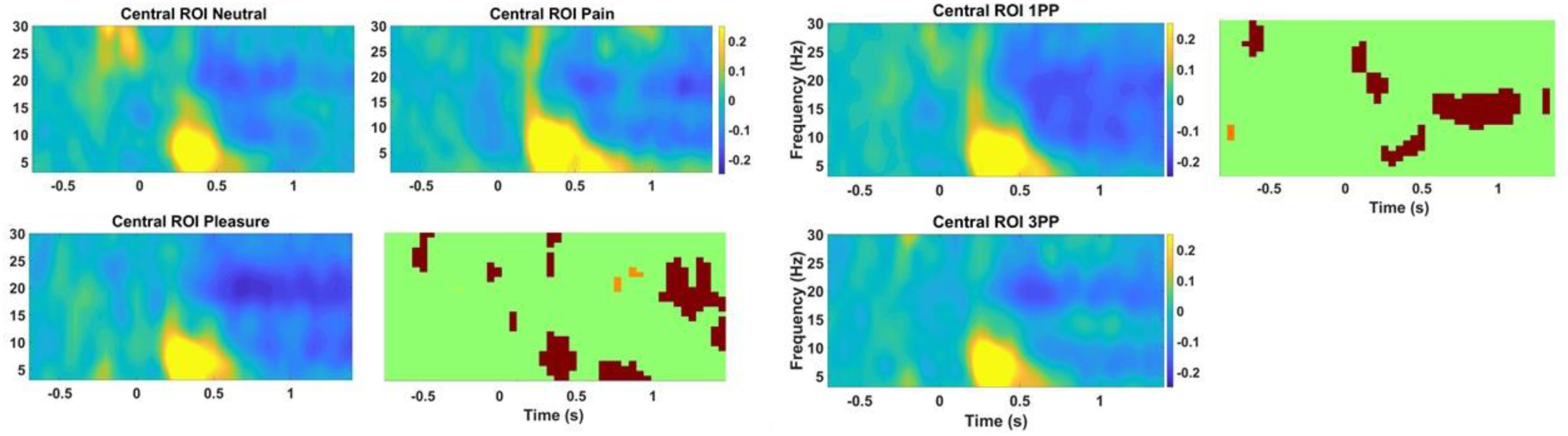
Time-frequency results. Main effects of Valence and Perspective Grand average of EEG total power for the two main effects of valence and perspective at the Central ROI (FCz, Cz, C3) and significant comparisons with corrected p-values within the green-background boxes. For the main effect of valence, single comparisons between the three levels (neutral, pain and pleasure), did not showed significant differences after correcting p-value for multiple comparisons (500 permutation, see methods). The main effect of perspective shows increased theta power in 1PP compared to 3PP between 300-500ms, and decreased alpha power in 1PP compared to 3PP between 500-1000ms.

**Figure 7.**
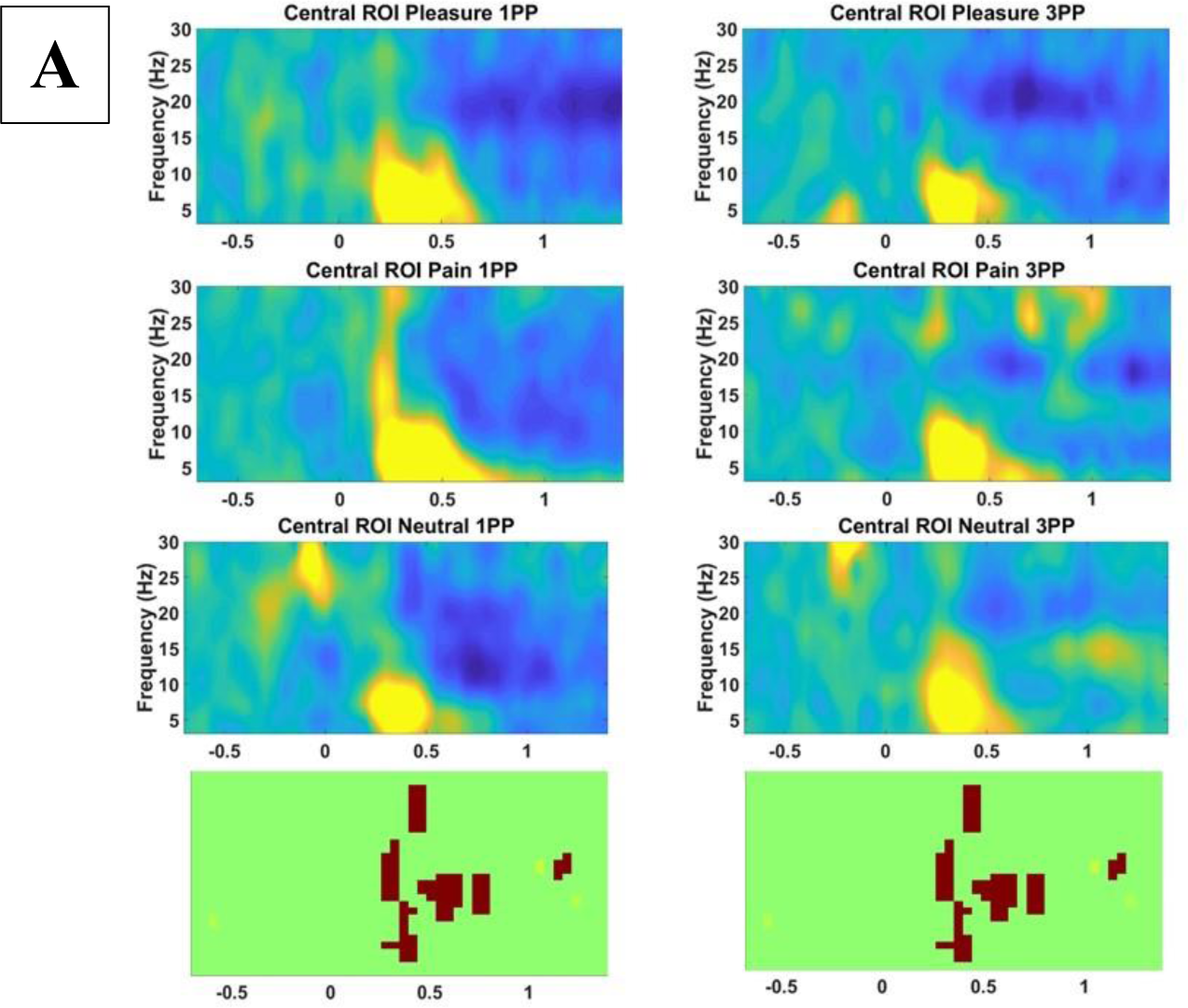
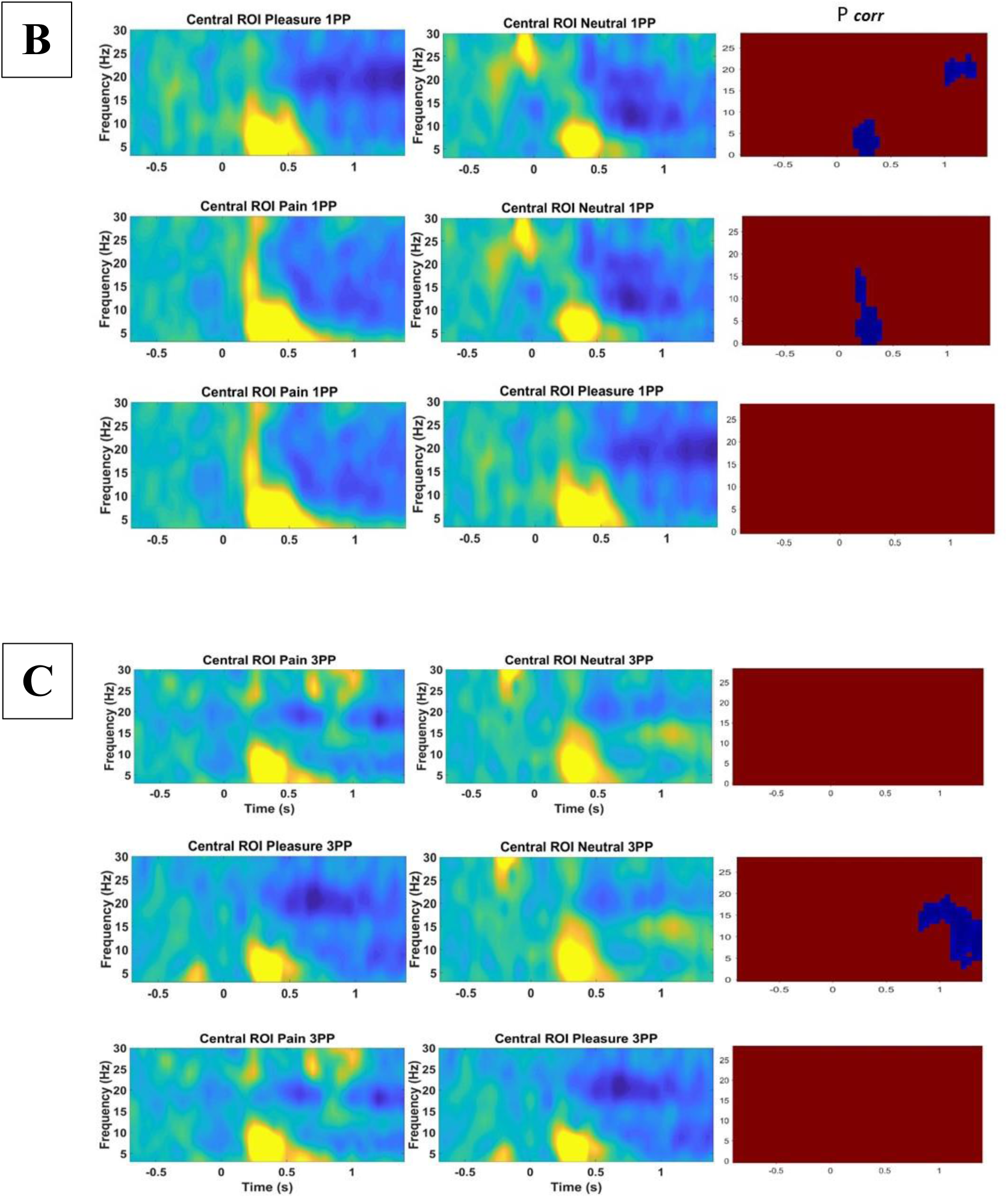
Time-frequency results. (A): Grand average of EEG total power at the central ROI and significant comparison within the green-background boxes for the significant interaction Valence x Perspective. (B) and (C): t-test for the significant interaction corrected for multiple comparisons. B is for 1PP and C for 3PP. Red-background boxes indicate significant results and corrected p-values

### Correlations between Subjective Ratings and EEG in the time-frequency domain

We used Pearson R correlation to test any possible association between the EEG power and the unpleasantness and intensity ratings. In specific, the analysis concerned subjective indices (computed subtracting the Neutral condition from either Pleasure or Pain) and time-frequency values that were significantly modulated in the different stimulus observation conditions. Ultralate Beta (1PP) and Theta (3PP) for the contrast between Pleasure and Neutral showed a moderate association with the Intensity ratings (r=0.38 and r=0.35, respectively). However, no association turned out to be statistically significant. All the ratings and across conditions comparisons are reported in Table 4.

**Table 4.** Correlations between Subjective Ratings and EEG in the time-frequency domain. We used Pearson correlation to test the associations between the EEG power and the Unpleasantness and Intensity ratings. Indexes were calculated (separately for each perspective) by subtracting the Neutral condition from Pleasure and Pain.

| Index | Frequency Band | Subjective Rating | Pearson R | p-value |
| --- | --- | --- | --- | --- |
| Pleasure 1PP - Neutral 1PP | Beta 800-1500ms | Pleasantness | 0.15 | 0.47 |
| Pleasure 1PP - Neutral 1PP | Beta 800-1500ms | Intensity | 0.38 | 0.07 |
| Pleasure 3PP - Neutral 3PP | Beta 800-1500ms | Pleasantness | 0.07 | 0.75 |
| Pleasure 3PP - Neutral 3PP | Beta 800-1500ms | Intensity | -0.18 | 0.40 |
| Pleasure 3PP - Neutral 3PP | Theta 800-1500ms | Pleasantness | -0.07 | 0.75 |
| Pleasure 3PP - Neutral 3PP | Theta 800-1500ms | Intensity | 0.36 | 0.08 |
| Pleasure 3PP - Neutral 3PP | Alpha 800-1500ms | Pleasantness | -0.08 | 0.70 |
| Pleasure 3PP - Neutral 3PP | Alpha 800-1500ms | Intensity | 0.12 | 0.57 |
| Pleasure 1PP - Neutral 1PP | Theta 200-350ms | Pleasantness | 0.09 | 0.66 |
| Pleasure 1PP - Neutral 1PP | Theta 200-350ms | Intensity | 0.01 | 0.97 |
| Pain 1PP - Neutral 1PP | Theta 200-350ms | Pleasantness | 0.26 | 0.22 |
| Pain 1PP - Neutral 1PP | Theta 200-350ms | Intensity | 0.15 | 0.50 |

## Discussion

In the present study we explored, for the first time using a within-subject design, the behavioural and brain reactivity to the observation of pain and pleasure stimuli delivered to an embodied vs. a non-embodied virtual avatar. To achieve our aim, we combined immersive virtual reality with time- and time-frequency domain EEG recording in healthy participants observing from a 1PP or 3PP: i) a virtual arm holding a needle that approached and then penetrated the right hand of an avatar (pain condition); ii) a virtual hand approaching the avatar’s right hand and caressing it (pleasure) and; iii) a virtual green ball approaching and touching the avatar’s right hand (neutral). Subjective and neurophysiological data indicate that cues to virtual embodiment like body ownership may operate at both explicit (Fusco et al., 2020; Monti et al., 2020; Tieri et al., 2015a) and implicit levels (Tieri et al., 2015b, 2017; Fusaro et al., 2016, 2021; Pavone et al., 2016; Casula et al., 2022; Pezzetta et al., 2023). Expanding on previous evidence (Tieri et al., 2015a; Pavone et al. 2016; Fusaro et al. 2016, 2019), subjective reports indicate that our experimental approach is adept to induce vicarious sensations and embodiment over artificial agents. Specifically, participants reported stronger illusory feelings of ownership over the virtual hand when observing it from a 1PP compared to a 3PP. Subjective ratings also indicate that our painful and pleasant virtual stimuli were effective in eliciting most unpleasant and most pleasant sensations, respectively, and were rated as more intense compared to neutral stimuli. These data confirm that IVR paradigms effectively reproduce veridical real-life like vicarious experiences. Tellingly, additional novel results of the present study concern the effect of affective salience, i.e. the effect of stimulus valence (painful, pleasurable, neutral) and the effect of observational perspective (1PP vs 3PP) on time- and time-frequency domain EEG findings.

### EEG results in the time domain

Valence and perspective differentially modulated early and late stimulus-evoked EEG components. In specific, early N2-VEPs and late EPN showed greater amplitude for pain and pleasure (vs. neutral), and for 1PP (vs 3PP). This pattern of results likely reflects attentional engagement, which turns out to be higher for emotionally valued stimuli compared with the neutral ones and for 1PP compared with the 3PP. N2-VEPs components are thought to reflect processes involved in selective attention (Olofsson et al., 2008; Luck and Kappenman 2011). Here, the early component did not distinguish between positive or negative value of the stimuli but coded its salience, either attentional or emotional (Todd et al., 2012). A greater EPN amplitude was found for 1PP compared to 3PP, and for Pain and Pleasure compared to Neutral conditions suggesting that this EEG component is linked to emotionally salient scenarios with no differences between positive and negative valence. Indeed, the EPN reflects automatic emotion-related influences on information (e.g., Olofsson et al., 2008; Schupp et al., 2004), processing of body parts (Hietanen et al., 2014), and processing of salient affective stimuli, with no category-specific sensitivity (Fabi & Leuthold, 2017).

Interestingly, vicarious pleasant touch elicited smaller LPP (400-600 ms) amplitude than painful and neutral stimuli. This result allows us to rule out that electrophysiological differences may be simply due to visual differences in the different scenarios, with pain and pleasure clips showing a syringe-holding or a touching virtual hand approaching the still hand and neutral ones only a ball. LPP has been proposed to reflect the arousal and the motivational salience of the stimulus (Cuthbert et al., 2000; Luck and Kappenman 2012; Hajcak et al., 2009), with greater amplitude for more relevant and arousing stimuli. However, studies indicate that central LPP can be modulated by the pleasantness of the received touch (Haggarty et al., 2020), with smaller LPP for more pleasant touch, in those people who preferred slow over fast touch (Schirmer et al., 2023). Thus, our result that LPP specifically responded to the pleasant scenario is in line with previous studies concerning real touch. Given the adopted paradigm, we cannot disambiguate if our LPP modulation was due to the pleasantness of the vicarious touch, or to its positive valence with respect to the other scenarios. Yet, our results expand current knowledge by showing that the effect of virtual pleasant touch is similar to what reported for real stimuli.

### EEG results in the time-frequency domain

EEG power in theta and alpha band was modulated by stimulus valence and perspective between 200-500 ms. Within this time-interval, for the theta band, the effect of valence led to an increase in power for pain and pleasure compared to neutral, while an increased power was found for 1PP scenarios compared to 3PP. The significant interaction scenarios X perspective interaction is explained by the modulation of theta power for pain and pleasure only in 1PP. The modulation of the theta-band frequency activity observed for affective stimuli is thought to be related to the emotional processing and affective valence discrimination, with enhanced theta power for affective stimuli, particularly in central and frontal areas (Aftanas et al., 2001; Mu et al., 2008; Schirmer et al., 2018). However, theta power increase has also been associated with cognitive effort (Gregory et al., 2022), sensorimotor integration (Kober & Neuper, 2011) and conflict/error-processing (Cavanagh et al., 2012; Pezzetta et al., 2018, 2023). Thus, the enhanced theta power for pain and pleasure vs neutral scenarios found in our study may be explained by the high cognitive load related to affective content. Moreover, finding enhanced theta-power only in 1PP suggests that different mechanisms may underpin vicarious experiences depending on the adopted perspective. Alpha band power was modulated by perspective only, showing power decrement in 1PP compared to 3PP, with no differences for the stimulus valence, nor an interaction between valence and perspective. Alpha band responses to emotionally arousing stimuli led to conflicting results, with pleasant and unpleasant stimuli associated with both alpha power increase or decrease respectively (Schubring & Schupp, 2019; Uusberg et al., 2013). However, less is known about the self-other distinction when stimuli are presented in ‘own’ vs. ‘other’ perspective. Alpha bands responses have been related to the allocation of visuospatial attention (Mathewson et al., 2011), the activation of sensory (Van Ede et al., 2011) and/or motor cortex (Perry et al., 2011), with alpha/mu suppression linked to social information processing (Pineda & Hecht, 2009) and perception of others’ pain (Cheng et al., 2008). Moreover, alpha suppression has been related to participants’ engagement in a task, with larger mu suppression when participants are engaged in a social game compared to passively looking at the other’s action (Oberman et al., 2007; Perry et al., 2011). In this vein, greater alpha suppression is associated with higher entrainment in time during a musical joint action paradigm (Novembre et al., 2016). Here, we found that the cognitive perspective rather than the stimulus valence, modulated alpha power in response to virtual scenarios, likely indicating different attentional engagement elicited by 1PP or 3PP respectively.

Finally, the significant interaction between valence and perspective also showed a power decrease after 800 ms, in the theta, alpha and beta band, an effect that was specific for pleasure scenarios. Interestingly, while theta power increase over midfrontal areas has been associated with conflict/error processing and valence discrimination (Aftanas et al., 2001; Cavanagh et al., 2012), theta power decrease in virtual reality has been associated with calmness (Eswaran et al., 2018). Beta desynchronization has been associated with motor readiness and sensorimotor activation (Riečanský et al., 2014; Fabi and Leuthold, 2017), the observation of a moving hand (Pfurtscheller et al., 2007), as well as cognitive aspects of pain processing (Valentini et al., 2017b). Moreover, while the power decrease for theta and alpha concerns only the 3PP, the power decrease for beta band is found for both perspectives. Overall, this power decrease seems specifically associated to pleasant stimuli and occurs in the ultra late time window. This effect may be associated with attentional engagement and sensorimotor integration contingent upon vicarious pleasant touch processing. Thus, although somewhat speculatively, we submit that, our decreased beta power may reflect a reduced alert or negative state. A similar meaning may be attributed to the the late time-window LPP amplitude reduction we observed for pleasure compared with neutral scenarios. Our findings of late and ultralate time- and time-frequency EEG signs of vicarious processing, is in keeping with a recent multivariate pattern analysis whole-brain EEG study showing that seeing touch evokes overlapping representations at later stages of information processing (Smit et al., 2023).

All in all, our study indicates that stimulus valence and perspective differentially affect ERPs related to attentional and motivational/affective processing, as well as EEG responses in theta, alpha and beta band. In particular, the affective scenarios (pain and pleasure) elicited similar EEG activity related to attentional engagement (N2, alpha and theta before 500 ms), while exhibiting distinct activity patterns at late and ultra-late time-window. Although these novel effects need further characterization, our patterns of results is compatible with the idea that vicarious pleasant experience, and its calming/consoling effect (Peled-Avron et al., 2018; Morrison, 2016) could be related to a specific cortical reactivity pattern, occurring after 500 ms in the neural processing. It is worth noting that no significant correlation between subjective reports and indices of neural activity is found. While we acknowledge that this is a possible limitation of the study, we remark that such lack of relation between subjective and objective data aligns with previous research using a similar approach and may be attributed to the dimension of the sample size that may not be sensitive enough for detecting subtle effects. A further possible limitation of the study is that the three clips, like other naturalistic complex stimuli used in neuroscience research, may differ in conspicuous visual or conceptual features. Studies indicate that seeing a hand vs a tool during visually guided actions may change electrocortical activity in the time-frequency domain (Mathie et al., 2023). Our pain and pleasure clips depict a virtual moving hand in addition to the static one, while the neutral clips depicts a ball and a static virtual hand. These differences may affect the electrocortical responses and make our interpretation ambiguous. However, most of our effects are related to the perspective, a condition where the virtual scenario for each clip is similar. More importantly, some of the effects concerning LPP are specific for pleasure, with pain and neutral stimuli showing similar patterns. This suggests that higher-order rather than comparatevely low-level factors modulate our reactivity to self vs other related, positive vs negative feelings.

## Conclusions

Our combined EEG and virtual reality study of how perspective-taking affects vicarious pain shows: i) a stronger salience of pain and pleasure with respect to neutral stimuli, particularly in 1PP, where the embodiment of the virtual avatar is maximal; ii) a reduction of the amplitude of a late positivity potential specifically associated with the observation of pleasure stimuli was also found. Our findings cast new light on the influence that perspective-taking exerts on the modulation of vicarious processing at behavioral and brain levels and have potential implications for clinical research, for example, in patients with abnormal perception or emotional reactivity to painful or pleasant stimuli (Boheme et al., 2020; Habig et al., 2021; Fusaro et al., 2023; Nicolardi et al., 2023).

## Acknowledgements

This work was supported by BIAL Foundation (n° 218/2016) and by Italian Ministry of Health (Ricerca Finalizzata, Giovani Ricercatori 2019, n°GR-2019-12369761). S.M.A. gratefully acknowledges the support of the Institut d’études avancées de Paris.

## Data availability statement

Raw data and scripts for the analyses will be available on a public data repository at the moment of publication of the manuscript in a peer-reviewed journal.

## Author contributions

G.T., M.F., V.N. and S.M.A. designed research; G.T. created the 3D models, software and implemented the VR setup; V.N., M.P.L. and M.M. collected the data; V.N. analyzed EEG data; G.T. and M.P.L analyzed subjective data; V.N. provided the first draft of the manuscript. S.M.A supervised the research. All authors approved the final version of the manuscript.

